# Relating C-reactive Protein to Psychopathology after Cardiac Surgery and Intensive Care Unit Admission: A Mendelian Randomization Study

**DOI:** 10.1101/196790

**Authors:** Lotte Kok, Bochao Danae Lin, Juliette Broersen, Erwin Bekema, Jelena Medic, Kristel R van Eijk, Manon H Hillegers, Dieuwke S Veldhuijzen, Jurjen J Luykx

**Author notes:** Corresponding author Lotte Kok, Department of Intensive Care Medicine, University Medical Center Utrecht, Utrecht, the Netherlands, Heidelberglaan 100, 3584 CX Utrecht, the Netherlands.

## Abstract

Patients admitted to an intensive care unit (ICU) are subjected to a high burden of stress, rendering them prone to develop stress-related psychopathology. Dysregulation of inflammation and, more specifically, upregulation of inflammatory markers such as C-reactive protein (CRP) is potentially key in development of post-ICU psychopathology.

To investigate the effects of state-independent CRP on symptoms of post-traumatic stress disorder (PTSD) and depression after ICU admission, we analysed the three leading single nucleotide polymorphisms (SNPs) of loci most strongly associated with blood CRP levels (i.e. rs2794520, rs4420638, and rs1183910) in an ICU survivor cohort. Genetic association was estimated by linear and logistical regression models of individual SNPs and genetic risk score (GRS) profiling. Mendelian Randomization (MR) was used to investigate potential causal relationships.

Single-SNP analyses were non-significant for both quantitative and binary trait analyses after correction for multiple testing. In addition, GRS results were non-significant and explained little variance in psychopathology. Moreover, MR analysis did not reveal any causality and MR-Egger regression showed no evidence of pleiotropic effects (p-pleiotropy >0.05). Furthermore, estimation of causality between these loci and other psychiatric disorders was similarly non-significant.

In conclusion, by applying a range of statistical models we demonstrate that the strongest plasma CRP-influencing genetic loci are not associated with post-ICU PTSD and depressive symptoms. Our findings add to an expanding body of literature on the absence of associations between trait CRP and neuropsychiatric phenotypes.

## Introduction

In the Netherlands, over 80,000 patients are admitted to the Intensive Care Unit (ICU) each year, and this number is increasing (https://www.stichting-nice.nl/). After discharge, health-related quality of life is largely determined by mental wellbeing (Iwashyna, 2010; Oeyen *et al,* 2010). The increasing number of ICU survivors combined with a decrease in quality of life renders in-depth studies into determinants of mental health in this patient category timely.

Post-traumatic stress disorder (PTSD) and depression are prevalent mental disorders following ICU admission (Asimakopoulou & Madianos, 2014). Several studies have demonstrated associations between acute high stress exposure due to ICU admission on the one hand, and PTSD and depression on the other (Davydow *et al,* 2013; Schelling *et al,* 2003). Despite the burgeoning body of research highlighting increased incidences of PTSD and depressive symptoms in ICU survivors (Davydow *et al,* 2008; Davydow *et al,* 2009), specific risk factors for developing post-ICU psychopathology remain largely undetermined. A better understanding of the potential mechanisms underlying the development of psychopathology after ICU admission could help to identify patients at risk of psychopathology following ICU stay.Since dysregulated inflammatory cascades are present in a high number of ICU patients, these might influence the development of psychopathology after discharge (Dieleman *et al,* 2012; Iwashyna, 2010). Evidence suggests that inflammation contributes to the development of PTSD and depression (Miller *et al,* 2009; Raison & Miller, 2011), and – in particular – increased C-reactive protein (CRP) levels are reported to be associated with posttraumatic stress disorder (PTSD), depression, and psychological distress (Danner *et al,* 2003; Eraly *et al,* 2014; Wium-Andersen *et al,* 2013).

Plasma CRP levels are highly affected by myriad state-dependent factors such as diet (King *et al,* 2003), physical activity (Kasapis & Thompson, 2004) and transient systemic infections. To circumvent the influence of these confounding factors on the traits of interest,we applied Mendelian Randomization (MR), a design that enables assessment of causality between a risk factor and an outcome. MR is based on the assumption that state-dependent features and environmental factors are generally equally distributed over carriers and non-carriers of genetic variants (de Haan *et al,* 2014; van der Weele *et al,* 2014). Thus, by taking genetic variants as proxies for circulating CRP (trait CRP), reverse causation and confounding are less likely to occur than when plasma CRP levels are considered (de Haan *et al,* 2014).

A genome wide association study (GWAS) including over 80,000 individuals found three single nucleotide polymorphisms (SNPs) – rs2794520, rs4420638 and rs1183910 – to be strongly associated with circulating CRP levels (Dehghan *et al,* 2011). The corresponding loci encompass the genes *CRP*, apolipoprotein C1 (*APOC1*) and hepatic nuclear factor 1-α (*HNF1A*), respectively (Dehghan *et al,* 2011). Although other CRP-related SNPs have been investigated in the context of PTSD or depressive symptoms (Ancelin *et al,* 2015; Halder *et al,* 2010; Michopoulos *et al,* 2015; Prins *et al,* 2016), the role that the abovementioned genetic variants play in ICU-related psychopathology have not been studied so far.

Here, we applied MR (including genetic risk scoring) to study the impact of trait CRP on psychopathology following ICU stay. We also investigated genetic pleiotropy between trait CRP and psychopathology following ICU stay. Our design has enabled us to assess causality, which to our knowledge had not been investigated to date.

## Methods

### Study design, participants and outcome measures

Data were drawn from the Dexamethasone for Cardiac Surgery study (DECS), a longitudinal follow up of patients from a multicenter, double-blind randomized controlled trial (Dieleman *et al,* 2012; Clinicaltrials.gov identifier NCT00293592, N=4494 participants). The aim of this trial was to compare high-dose intravenous dexamethasone (1 mg/kg bodyweight, not exceeding 100 mg) with placebo administration in patients undergoing cardiac surgery who were subsequently submitted to the ICU. Eight cardiac surgery centers in The Netherlands participated, and the study was approved by all applicable research ethics committees (Dieleman *et al,* 2012). The medical Ethics Committee of the University Medical Center Utrecht approved the current follow-up protocol. All subjects participating in the current study signed both the initial and additional informed consent forms.

Participants were recruited between April 13, 2006 and November 23, 2011. DECS included 4494 randomized adult patients requiring a cardiopulmonary bypass during cardiac surgery. Of the original DECS study population, 2458 patients (recruited from five participating centers) were eligible for the follow-up study as the remainder had passed away or exceeded the inclusion criterion of maximum follow up time (i.e., 48 months after randomization).

PTSD symptoms were measured using the Self-Rating Inventory for PTSD (SRIP), a validated Dutch questionnaire (Hovens *et al,* 2002). This 22 item questionnaire addresses DSM-IV-TR criteria of PTSD in the past four weeks. The validated Beck Depression Inventory (BDI-II) (Beck *et al,* 1998) was used to measure depressive symptoms. Items on both questionnaires are scored on a 0-3 Likert scale, with higher scores indicating more severe symptoms. For binary trait analyses, we rated symptoms of PTSD as present in the event of SRIP scores above the validated threshold of 39, which was the case in 10.8% of all participants, comparable to what has been described in previous studies (van Zelst *et al,* 2003). Symptoms of depression were present in 13.2% of all participants, after applying threshold of 13.5 to the BDI-II score (Dozois *et al,*1998). A detailed description of the patient selection and measurements is reported elsewhere (Kok *et al,* 2016). For the analyses in this study, we log transformed SRIP and BDI-II variables due to non-normal distributions.

### Genotyping procedures

Genomic DNA was extracted from saliva using salivettes (Kleargene™ XL nucleic extraction kits) according to the manufacturer's protocol. Quantity of DNA was measured using QuantiT™ PicoGreen®dsDNA reagent fluorescence by Varioskan™ Flash Multimode reader (Thermo Scientific™). Genotypes were determined by real-time polymerase chain reaction (PCR) conducted in QuantStudio™ 6 (Applied Biosystems by Life Technologies™) with Custom TaqMan® SNP Genotyping Assays for rs27924520 and rs1183910, and Assays-by-Design^SM^ for rs4420638 (Applied Biosystems by Life Technologies™). For these analyses a high-throughput dried-down DNA method was used. Herein, 2.5 µL Master Mix (2X, TaqMan Genotyping Master Mix, Applied Biosystems by Life Technologies™) and 0.125 μL of a specific assay (TaqMan Genotyping 40X Assay, Applied Biosystems by Life Technologies™) and 2.375 μL Milli-Q (H2O) per sample were added. The real-time PCR procedure included three stages: pre-read fluorescence measurement (30 seconds) at 60.0°C, 40 two-step cycles: DNA denaturation at 95.0°C (15 seconds) and amplification of SNP sequence of interest at 60.0°C (60 seconds) and post-read fluorescence measurement (30 seconds) at 60.0°C. Allelic discrimination was determined by the post-read stage fluorescence intensity using QuantStudio™ 6 genotyping software.

## Statistical analyses

Descriptive statistics were generated using Statistical Package for Social Sciences (SPSS Inc., version 20.0. IBM corp. Chicago, IL, USA), while genetic analyses were performed using PLINK v1.07 (Purcell *et al,* 2007) and R (www.r-project.org). Missing values at baseline and psychopathology outcome variables were imputed using multiple imputation in SPSS. Ten imputed datasets were created based on independent and dependent variables. Analyses were conducted in each dataset and results were pooled according to Rubin’s rule (Rubin, 1987). The Hardy-Weinberg equilibrium (HWE) threshold was set at *p*>0.05. To comprehensively examine both single-SNP and cumulative effects of all three SNPs on psychopathology following ICU stay, the following analyses were carried out subsequently.

First, per-SNP linear regression models with SRIP and BDI-II scores as outcome variables were performed with the following covariates: sex, age, duration of ICU admission, treatment (dexamethasone vs. placebo administration) and genotype-treatment interaction. Treatment was included because of the potent anti-inflammatory properties of dexamethasone. The significance threshold was Bonferroni corrected (α=0.05/3; α=0.017) to account for the three SNPs tested.

Second, to model the cumulative effect of the three SNPs on PTSD and depression, we used genetic risk score (GRS) profiling (Purcell *et al,* 2009). Effect alleles were presented as “coded alleles” effect sizes were presented as beta-values and obtained from the CRP GWAS (N=80,000 individuals; Dehghan *et al,* 2011) to calculate GRS in PLINK. GRS was regressed against log transformed SRIP and BDI-II scores to calculate the proportion of explained variance in psychopathology outcome. Two models were applied: the first model only included sex and age as covariates, while the second model included all eight variables mentioned before as covariates.

Third, to corroborate the results of our GRS model, we performed a fixed-effect (inverse variance weighted, IVW) meta-analysis in line with a recent MR study (Vaucher *et al,* 2017) using the Mendelian Randomization package in R (Yavorska & Burgess, 2017). The two-sample MR IVW method reports estimates of causality between exposures and outcomes. All effect sizes were in standard deviation units for continuous traits and log odds ratios for disease and binary traits. The standard error (SE) was estimated using the delta method (Bautista *et al,* 2007; Thomas *et al,* 2007). Multiple sensitivity analyses validated the three MR assumptions (i.e., [1] the genetic marker is robustly associated with exposure, [2] the genetic marker is conditionally independent of outcome [pleiotropy], and [3] the genetic marker is conditionally independent of measured/unmeasured confounders), as well as the NO Measurement Error (NOME) assumption (Bowden *et al,* 2016). Moreover, the more robust MR-Egger method (Spring *et al,* 1989) was applied to investigate potential directional horizontal pleiotropy. The high I^2^ statistic (>90% in our study) supports the MR assumption of NOME. In addition, leave-one-out analysis was performed (Stone, 1974). In line with the abovementioned quantitative traits studies, per-SNP logistic regression to assess single-SNP effects was performed first, followed by logistic regression on GRS calculated from the three SNPs, and concluded with fixed-effects meta-analysis.

Finally, to assess whether effects of CRP-associated SNPs are specific to post-ICU admission psychopathology or also generalizable to 14 other psychiatric disorders, we used the MR-base package (Hemani *et al,* 2016), aiming to investigate any causal relationships between CRP and such phenotypes, based on the abovementioned three leading CRP-associated SNPs. If these candidate SNPs could not be extracted from the statistical data summary, we extracted a proxy SNP instead (the most informative SNP having highest LD r^2^-value). The significance threshold was again Bonferroni corrected (α=0.05/14=0.0036).

## Results

### Demographics and clinical characteristics

The study population consisted of 1093 subjects for whom signed informed consent, returned completed questionnaires and sufficient DNA were available (Figure S1). The median age of the participants was 69 years (interquartile range (IQR): 63-76) and ranged between 26 and 91 years. The majority of patients were male (78%). Dexamethasone was administered to 49.8 percent of the participants while 50.2 percent received placebo. Patients were admitted to the ICU for 5 to 1800 hours with a median duration of stay of 21 hours (IQR: 18.5-23.5). Demographic and clinical characteristics are displayed in Table 1.

**Table 1.** Demographic and clinical characteristics of the study population (N=1093).

| Characteristic | N (%) | Median (IQR) | Range |
| --- | --- | --- | --- |
| Demographics |  |  |  |
| Age (years) | - | 69.6 (63.1-76.1) | 26-91 |
| Sex – Male | 854 (78.1) | - | - |
| ICU |  |  |  |
| Dexamethasone treatment | 544 (49.8) | - | - |
| Duration of ICU stay (hours) | - | 21.0 (18.5-23.5) | 5-1800 |
| Self-reported psychiatric history |  |  |  |
| Depression | 22 (2.0) | - | - |
| Bipolar Disorder | 6 (0.5) | - | - |
| Anxiety Disorder | 17 (1.6) | - | - |
| Other Psychiatric Disorders | 13 (1.2) | - | - |
| Outcome measures (total scores) |  |  |  |
| SRIP |  | 26.0 (21.5-30.5) | 22-78 |
| BDI-II |  | 5.0 (1.5-8.5) | 0-51 |

### Genotyping success rates and Hardy-Weinberg equilibrium

The genotyping success rates for rs2794520, rs4420638 and rs1183910, were 97.7, 96.5, and 97.5 percent, respectively. All three SNPs were in Hardy-Weinberg equilibrium (HWE; p=1.0; p=0.076, and p=1.0, respectively) and the minor allele frequencies (MAFs) were 0.3455, 0.1909 and 0.2935, which is in agreement with the 1000 Genomes project (http://www.1000genomes.org). Genotype distributions are presented in Table S1.

### Mendelian randomization analyses

None of the three SNPs was individually associated with PTSD or depressive symptoms at the Bonferroni-corrected significance threshold (α=0.017; Table 2a and 2b). rs2794520 showed nominal evidence of association with quantitative PTSD symptoms (β: 0.2142, p=0.0325) and with quantitative depression symptoms (β: 0.2203, p=0.0273): the CRP-decreasing T-allele (which is also the minor allele) was associated with increased risk of psychopathology following ICU stay (Figure 1a and 1b). Logistic regression showed no significant results for the binary traits either: presence of the T-allele of rs2794520 tended to increase the risk of depression (OR=1.811, p=0.051) and PTSD (OR=1.409, p=0.316) non-significantly.

**Figure 1a.**
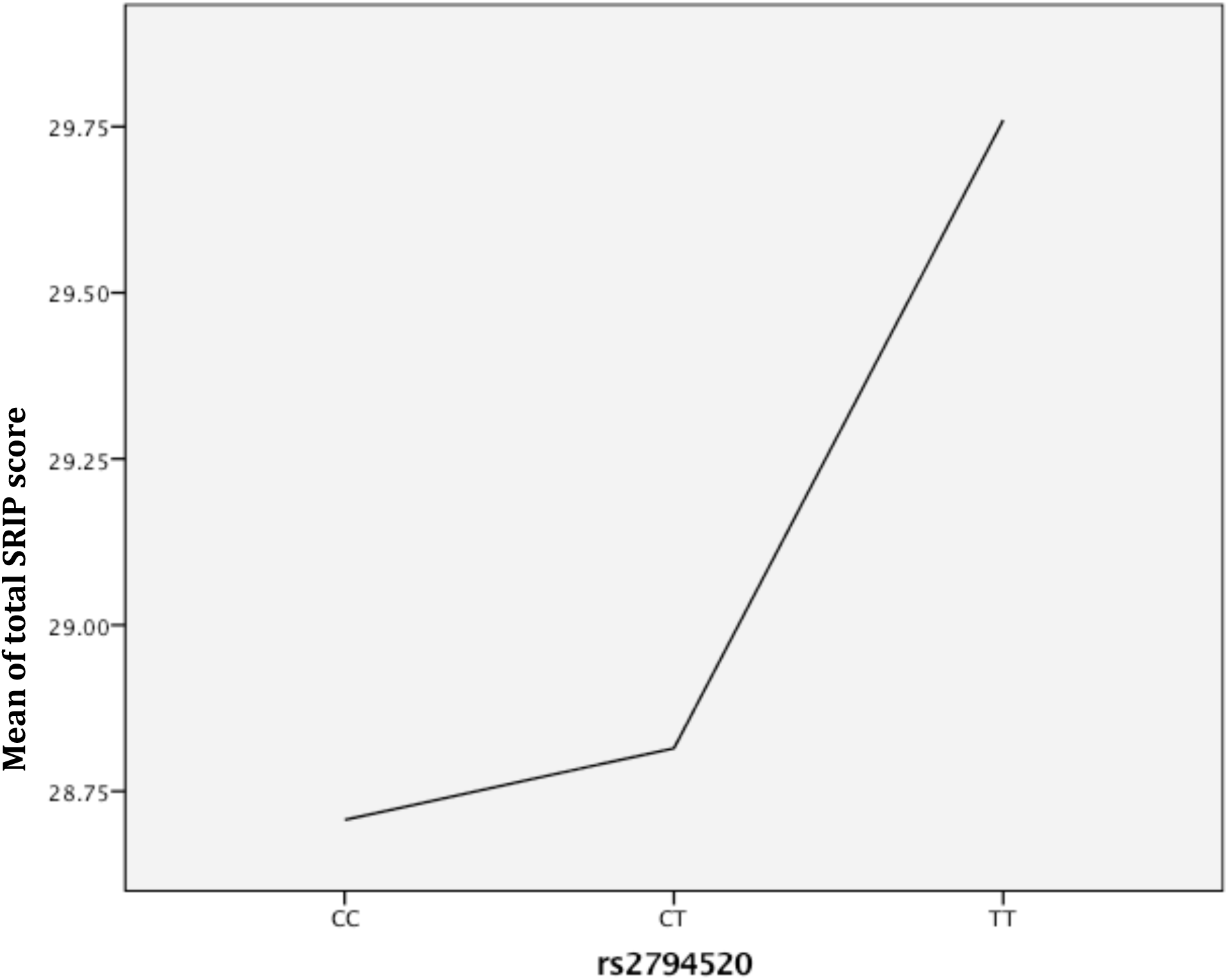
Means plot of PTSD symptoms per genotype.

**Figure 1b.**
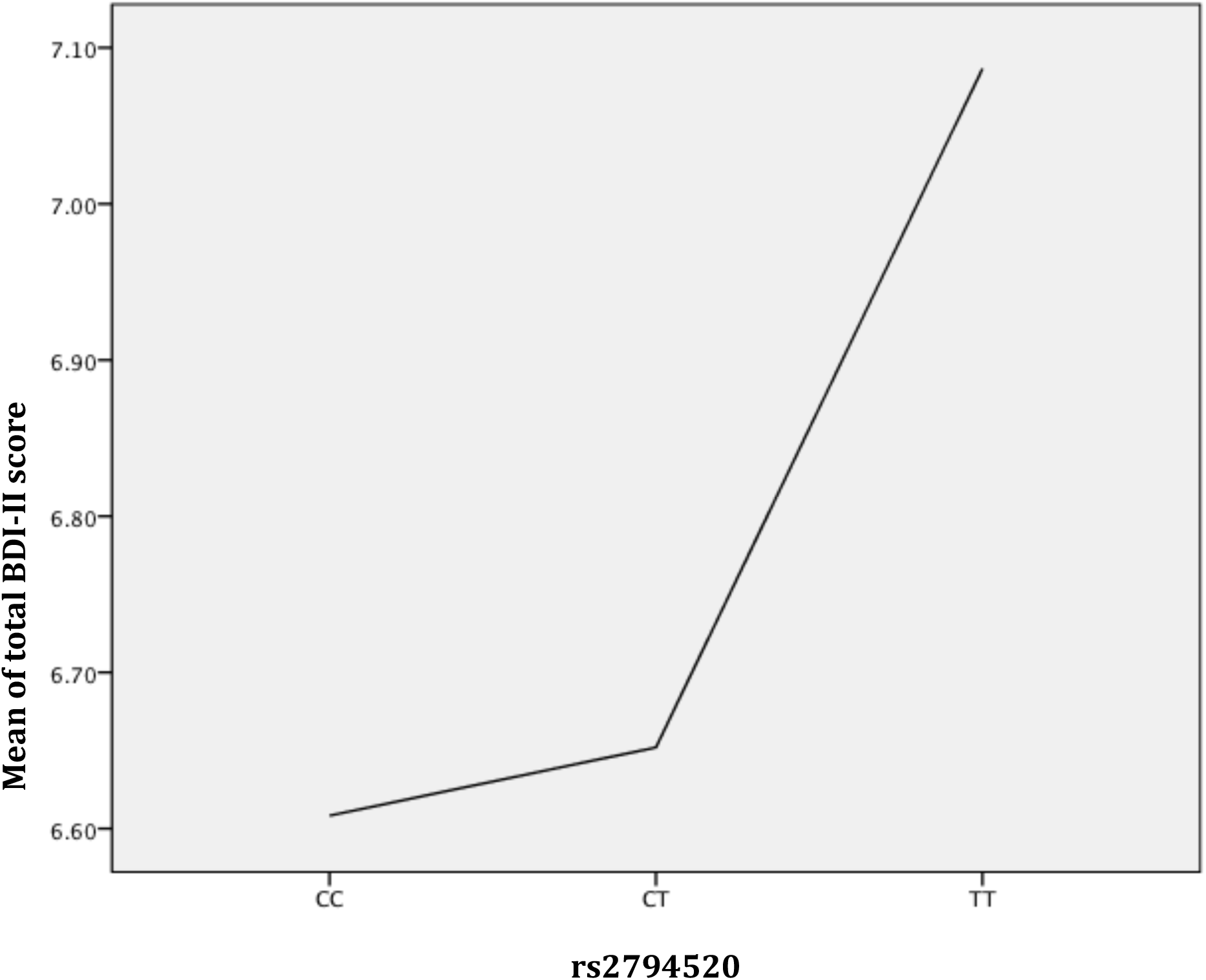
Means plot of depressive symptoms per genotype.

**Table 2a.**
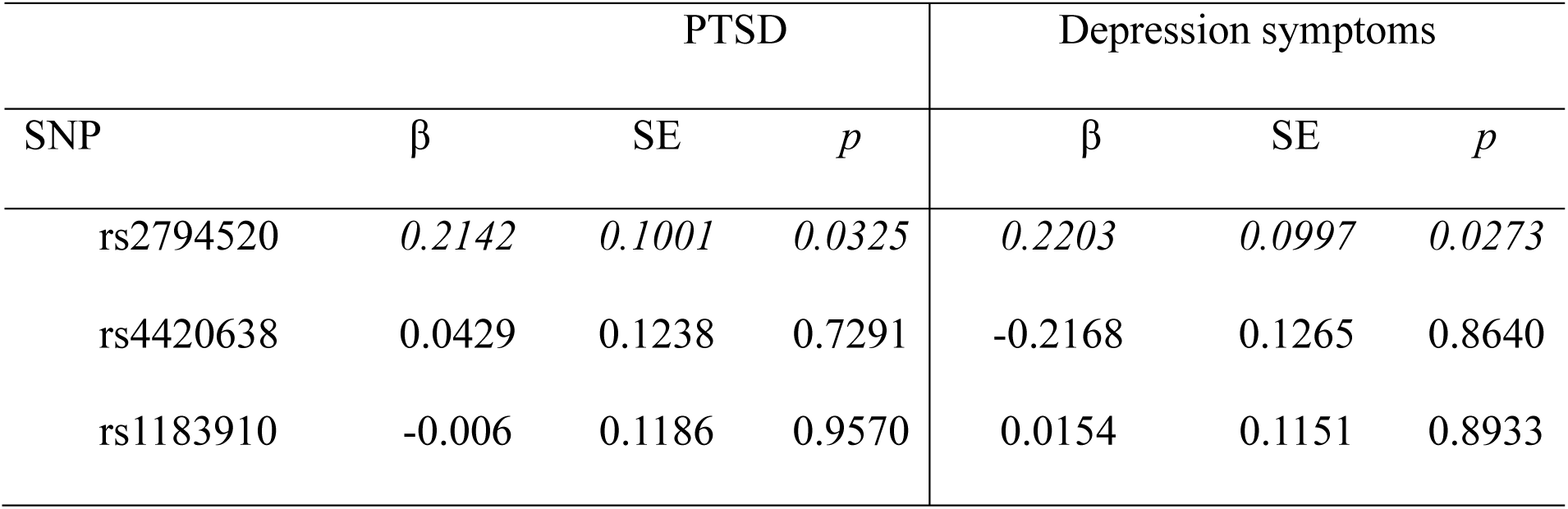
Association between CRP-associated SNPs and psychopathology symptoms: linear regression.

**Table 2b:**
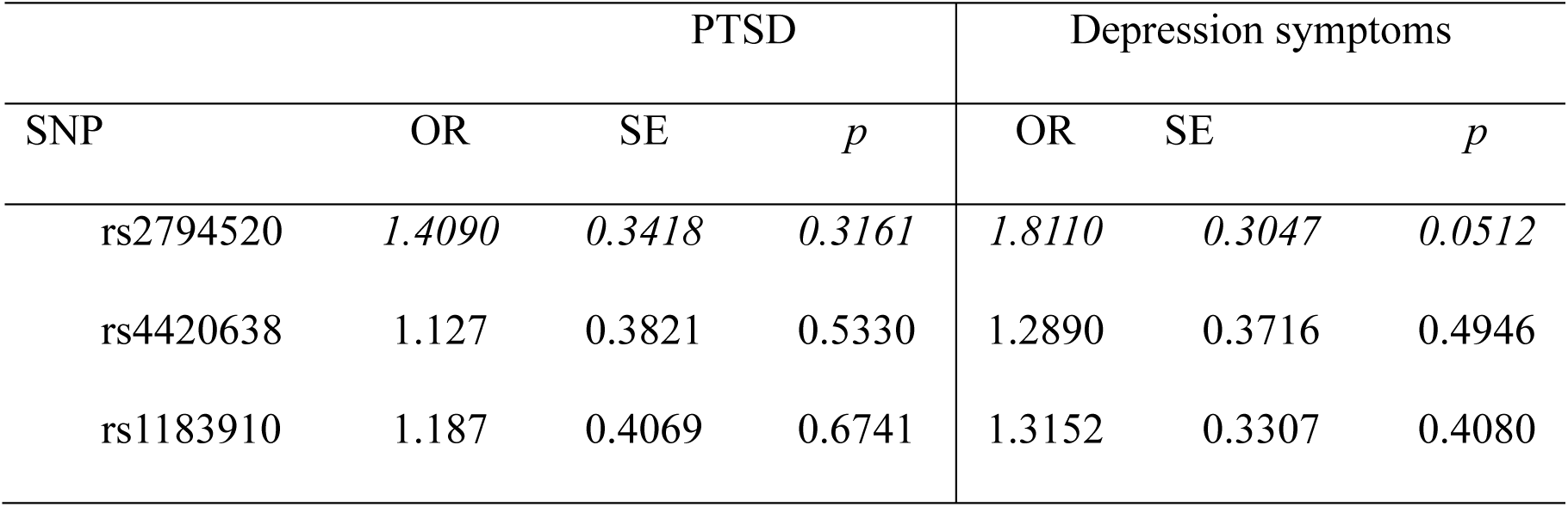
Association between CRP-associated SNPs and psychopathology symptoms: logistic regression.

When applying GRS, the proportions of variances of quantitative and binary PTSD and depressive symptoms explained by the CRP loci were non-significant (p>0.05) and below 0.0035 in both models (Figure S2).

Meta-analysis of the effects of the three CRP SNPs on PTSD and depressive symptoms was similarly negative (Figure S3, Figure S4, and Table S2a). There was no evidence of heterogeneity (p=0.02). MR-Egger regression (Table S2b) consistently showed no causal effects: the I^2^ statistic (indicating the level of instrument bias in the context of MREgger) was high (>90%). Therefore, the expected relative dilution did not affect the standard MR-Egger analyses for our data. Directional horizontal pleiotropy of CRP and psychopathological outcomes was absent (*p*-pleiotropy>0.05). Leave-one-out sensitivity analyses (Figure S5) showed none of the SNPs altering pooled effect sizes, indicating reliable and stable meta-analytic results. In line with this, parallel binary trait analysis showed no association between either individual SNPs or GRS on the one hand and psychopathology outcomes on the other, pleading against a causal relation between CRP-levels and psychopathology.

Finally, estimation of causality between the three CRP SNPs and 14 other psychiatric disorders by MR-base showed that level of CRP does not significantly affect the occurrence of other neuropsychiatric diseases (Table 3). No directional horizontal pleiotropy effects were detected among CRP and these neuropsychiatric traits.

**Table 3:** Two sample MR results from MR-base for CRP and various other neuropsychiatric disorders.

| Phenotypes | PMID | sample size | test statistics | OR | 95%CI | p-value | p-pleiotropy |
| --- | --- | --- | --- | --- | --- | --- | --- |
| Schizophrenia | 25056061 | 82,315 | log odds | 0.896 | 0.820-0.978 | 0.014 | 0.356 |
| Anorexia nervosa | 24514567 | 17,767 | log odds | 0.958 | 0.594-1.543 | 0.858 | 0.965 |
| Bulimia nervosa | 23568457 | 2,442 | log odds | 0.983 | 0.923-1.047 | 0.603 | 0.591 |
| Neuroticism | 25993607 | 160,958 | arbitrary<br>score | 0.983 | 0.907-1.066 | 0.685 | 0.761 |
| Alzheimer's disease | 24162737 | 54,162 | log odds | 0.239 | 0.006-8.482 | 0.432 | 0.122 |
| Bipolar disorder | 21926972 | 16,731 | log odds | 1.209 | 0.929-1.576 | 0.158 | 0.621 |
| Parkinson's disease | 19915575 | 5,691 | log odds | 0.651 | 0.459-0.924 | 0.017 | 0.355 |
| Autism | 23453885 | 29,415 | log odds | 1.091 | 0.832-1.430 | 0.528 | 0.624 |
|  | PGC 2015 | 10,263 | log odds | 1.077 | 0.896-1.349 | 0.548 | 0.560 |
| Major depressive disorder | 22472876 | 18,759 | log odds | 1.080 | 0.914-1.276 | 0.366 | 0.758 |
| Depressive symptoms | 27089181 | 161,460 | SD | 1.002 | 0.971-1.033 | 0.910 | 0.782 |
| Subjective well being | 27089181 | 298,420 | SD | 1.019 | 0.997-1.042 | 0.093 | 0.994 |
| Internalizing problems | 24839885 | 4,569 | NA | 0.973 | 0.814-1.162 | 0.759 | 0.563 |
| Amyotrophic lateral sclerosis | 27455348 | 36,052 | log odds | 1.014 | 0.986-1.043 | 0.339 | 0.373 |
| PTSD (European population) | 25904361 | more than 5,000 | log odds | 1.078 | 0.811-1.432 | 0.607 | 0.373 |

## Discussion

The present study was designed to examine the potential effect of trait CRP on PTSD and depressive symptoms in a large cohort of ICU survivors (N=1093). Overall, no evidence for a causal relationship between trait CRP and psychopathology after ICU-stay was found. In addition, we demonstrated that these traits are unlikely to share non-polygenic genetic overlap. Finally, we showed that the absence of causal effects of plasma CRP also applies to other neuropsychiatric traits. Since these disorders share many risk factors, which further pleads against a role of CRP in the development of psychopathology. Considering the many MR assumptions, negative MR findings are particularly valuable (van der Weele *et al,* 2014). Provided the genetic variants are strongly associated with the exposure – which is the case in this study – negative MR findings more strongly advance insights into relationships between exposures and traits than positive MR findings (van der Weele *et al,* 2014). Moreover, a null association (with a narrow confidence interval) between a strong instrument and an outcome provides more robust evidence of no or a very small causal effect.

Several factors may explain these findings. First, being admitted to an ICU is commonly perceived as stressful and this may trigger the development of PTSD and depressive symptoms (Schelling *et al,* 2003; Davydow *et al,* 2008). Thus, environmental factors (e.g. illness severity and duration of stay) may be stronger determinants of psychiatric outcomes following ICU stay than genetic underpinnings. Elevated levels of CRP in the context of psychopathology as detected in previous studies (Danner *et al,* 2003; Eraly *et al,* 2014; Wium-Andersen *et al,* 2014) may also occur due to other mechanisms, such as reversed causation, where PTSD and/or depressive symptoms cause elevated CRP levels rather than vice versa. Upregulation of inflammatory markers – e.g. state CRP – could therefore still be part of the pathophysiological pathway involved with the development of stress-related psychopathology (Duivis *et al,* 2013; von Känel *et al,* 2007). Future studies combining plasma CRP levels with MR could provide a more-refined understanding of the relationship between CRP levels and psychopathology (Almeida *et al,* 2009; Wium-Andersen *et al,* 2014). Furthermore, CRP-related genetic mechanisms other than single-locus effects – e.g., epistasis (for which this study was underpowered) and epigenetics – may influence the relationship between CRP and psychopathology. Finally, our power analysis indicates that our model had 80% power to detect an effect size of 0.007. This indicates that our study was adequately powered to detect medium to large effects, but underpowered to pick up very small effects (Freeman *et al,* 2013).

Although a positive association between CRP and psychopathology has been observed (Wium-Andersen *et al,* 2014; Ma *et al,* 2010), causal effects of CRP-associated loci on neuropsychiatric diseases had remained debatable. A recent MR study of CRP with multiple complex traits in a European study population, in which two CRP genetic risk scores as instrumental variables were used, showed no causality between CRP and autism, bipolar disorder, Parkinson’s disease, major depressive disorder, and a negative effect on schizophrenia (i.e., elevated levels of CRP are not predisposing but in fact protective) (Prins *et al,* 2016). Similarly, in our MR-based study, a suggestive but nevertheless consistently negative effect on schizophrenia was detected (p=0.014, which was the lowest p-value among 14 psychiatric traits) (Dehghan *et al,* 2011; Ripke *et al,* 2014). Although the negative effect on schizophrenia was not significant after correction for multiple testing, the validation of this result requires further investigation of more CRP-associated SNPs. No significant directional horizontal pleiotropy for the three CRP-associated SNPs was detected in a broad selection of psychiatric disorders (p-pleiotropy >0.05), implying these psychiatric disorders and state CRP levels are not associated with the CRP genetic variants via separate pathways.

Since other studies implicate different CRP polymorphisms from the ones we investigated, comparison of our results with previous findings is challenging. For example, rs1130864 (4kb away from rs2794520, r^2^=0.198) was found to be associated with more PTSD symptoms in African American population. With regard to depressive symptoms, a limited number of studies have been conducted, showing ambiguous results. Three studies found no association between these genetic variants on CRP genes and depression (Halder *et al,* 2010; Luciano *et al,* 2010; Wium-Andersen *et al,* 2014), while one found CRP polymorphisms rs1800947 and rs3093059 to be associated with depression irrespective of plasma CRP levels (Ancelin *et al,* 2015).

It is important to bear in mind that this study addressed a cardiac surgery cohort. In general, these are relatively healthy patients who are only admitted to the ICU overnight. Nevertheless, cardiovascular disease has been associated with inflammatory dysregulation in general (Bot & Kuiper, 2017). Differentiation between the effect of ICU treatment and genetic predisposition for the development of PTSD and depression is therefore challenging in our study population. Targeting phenotypically more diverse cohorts of critically ill patients may result in a more fine-grained understanding of the relationship between trait CRP and psychopathology. On the other hand, strengths of our study comprise the investigation of the most strongly associated CRP genetic loci, the high genotyping success rates and the stringent and comprehensive statistical analyses including several sensitivity analyses, while correcting for multiple testing and relevant covariates.

In conclusion, based on our MR method it is unlikely that trait CRP and psychopathology following ICU stay are causally related or demonstrate high degrees of genetic overlap. Identification and investigation of novel CRP loci and inflammatory signalling pathways may result in a better understanding of biological mechanisms involved in the relationship between inflammation and psychopathology following ICU stay. Furthermore, a deeper understanding of the relationship between state and trait inflammatory processes on the one hand, and psychopathology on the other, may in the long run spur identification of ICU patients at risk. Such knowledge in turn may open avenues for targeted preventative measures and treatments in these individuals at risk of developing psychopathology.

## Acknowledgements

The authors gratefully acknowledge the time and effort of the centers that participated in this study. We thank Diederik van Dijk and Marian Joëls for giving us the time and opportunity to perform this study.

## Figure Legends

**Figure 1a.** This graph depicts how subjects homozygous for the minor T-allele of rs2794520 seem vulnerable to develop PTSD symptoms and to promote the development of depressive symptoms after ICU stay. The y-axis represents mean total score of the SRIP questionnaire and thus severity of PTSD symptoms, whereas genotypes of rs2794520 are shown on the x-axis.

**Figure 1b:** This graph depicts how subjects homozygous for the minor T-allele of rs2794520 seem vulnerable to develop depressive symptoms and to promote the development of depression symptoms after ICU stay. The y-axis represents mean total score of the BDI-II questionnaire and thus severity of depressive symptoms, whereas genotypes of rs2794520 are shown on the x-axis.

**Figure S1**. ‘n’ represents numbers of patients included or excluded. Arrows display included patients, while lines present excluded patients.

**Figure S2.** R^2^: multiple R^2^ in linear regression, Naegelkerke’s pseudo R^2^ in logistical regression. Dichotomous trait for PTSD and depression using validated cut-off scores. Model 1 contains GRS, age, and sex as covariates. Model 2 contains GRS, age, sex, length of ICU stay, dexamethasone vs. placebo treatment and genotype-treatment interaction as covariates.

**Figure S3.** Forest plots of meta-analysis (fixed-effects, inverse variance) results for the three SNPs on PTSD and depressive symptoms after ICU stay.

**Figure S4.** snp1: rs4420638, snp2: rs2794520, snp3: rs1183910. SNP-exposure (associations with *CRP*) and SNP-outcome (associations with psychopathology outcomes) coefficients used in Mendelian Randomization analysis. Error bars (95% Confidence Interval) are reported for each association. The blue lines represent fix–effect meta-analysis (inverse variance weighted; IVW) model estimations.

**Figure S5.** Forest plots of leave-one-out sensitivity analysis (fixed-effects, inverse variance) results for the three SNPs on PTSD and depressive symptoms after ICU stay.

## Table Legends

**Table 1:** Values are presented as either ratios or medians with inter quartile ranges (IQR) and full ranges. BDI-II, Beck Depression Inventory II score; ICU, intensive care unit; SRIP, Self-Report Inventory for PTSD score.

**Table 2a:** Displayed are standardized beta-coefficients (β), standard error (SE) and p-values (*p*). The suggestively significant finding is in italics.

**Table 2b:** Displayed are standardized odds ratio (OR), standard error (SE) and p-values (*p*). The suggestively significant finding is in italics.

**Table 3.** Displayed are PubMed ID (PMID), sample size, test statistics used as instruments from GWAS studies for psychosocial disorders; odd ratio (OR), 95% confidence interval (95%CI), nominal p-values (p-value) from MR-base analyses with IVW models, and p-values for pleiotropy (p-pleiotropy) test from MR-base analyses with Egger model.

**Table S1.** Observed minor allele frequencies (MAFs) based on our study population. Expected MAFs are according to the 1000 Genomes project.

**Table S2a:** shown are the results for the meta-analysis (fixed-effects, inverse variance) for the three SNPs on PTSD and depressive symptoms after ICU stay.

**Table S2b:** shown are the results for the meta-analysis (MR-Egger regression) for the three SNPs on PTSD and depressive symptoms after ICU stay.

## Supplementary Material and Methods

**Figure S1.**
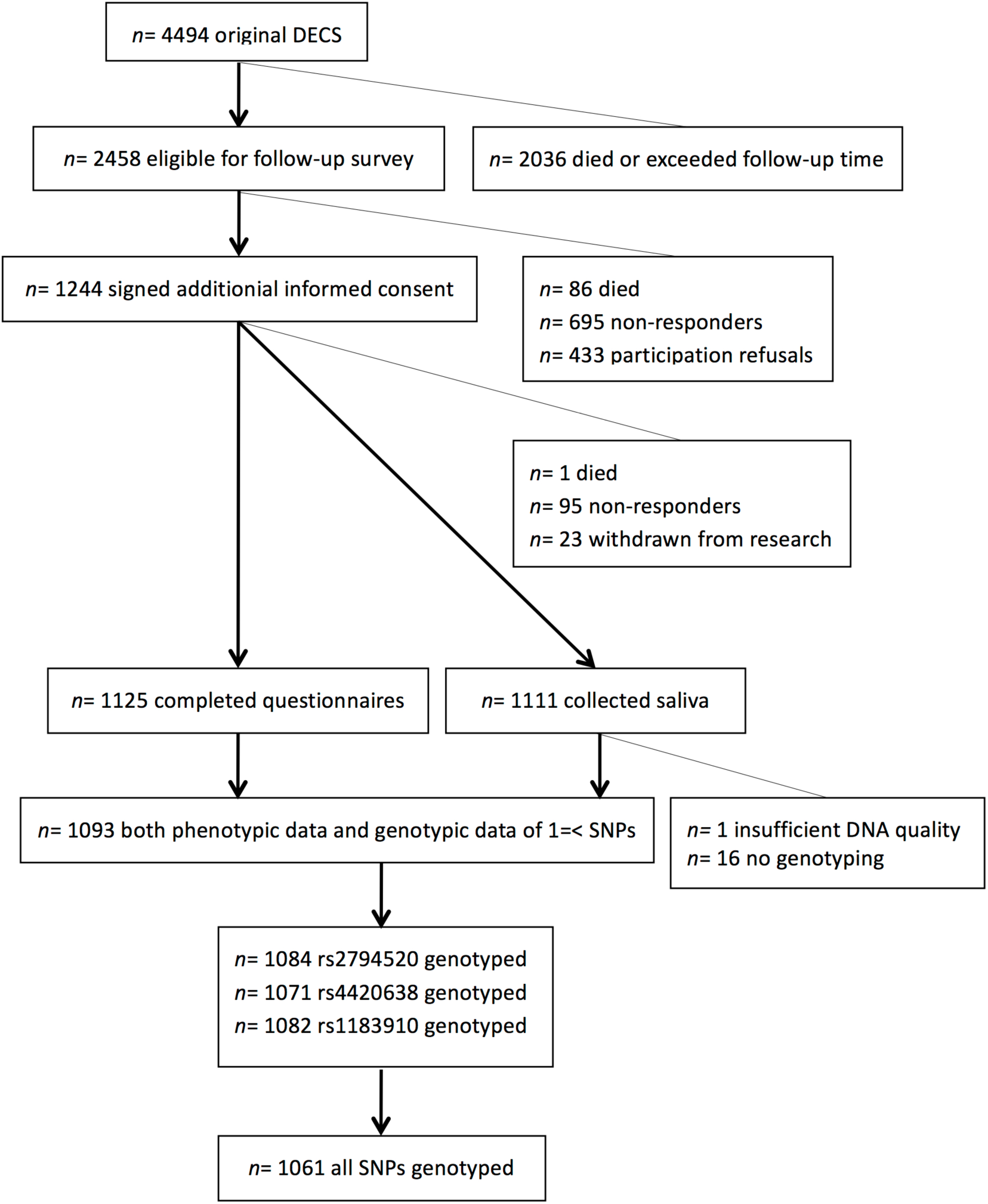
Flow chart of recruitment of participants.

**Figure S2.**
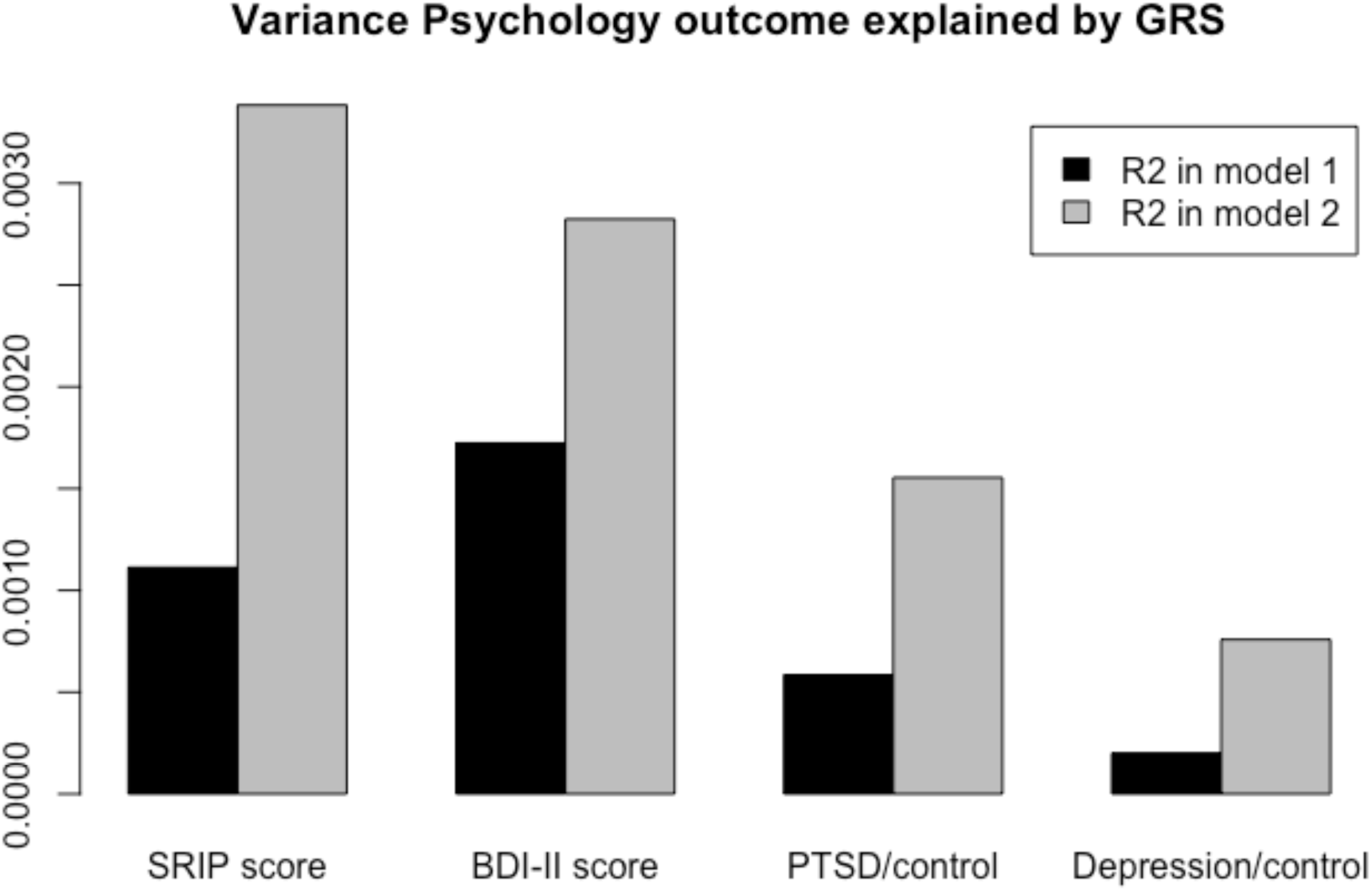
Proportion of variance in psychopathology explained by polygenic risk scores (GRS) obtained from the three CRP-associated SNPs using individual-level genetic data.

**Figure S3.**
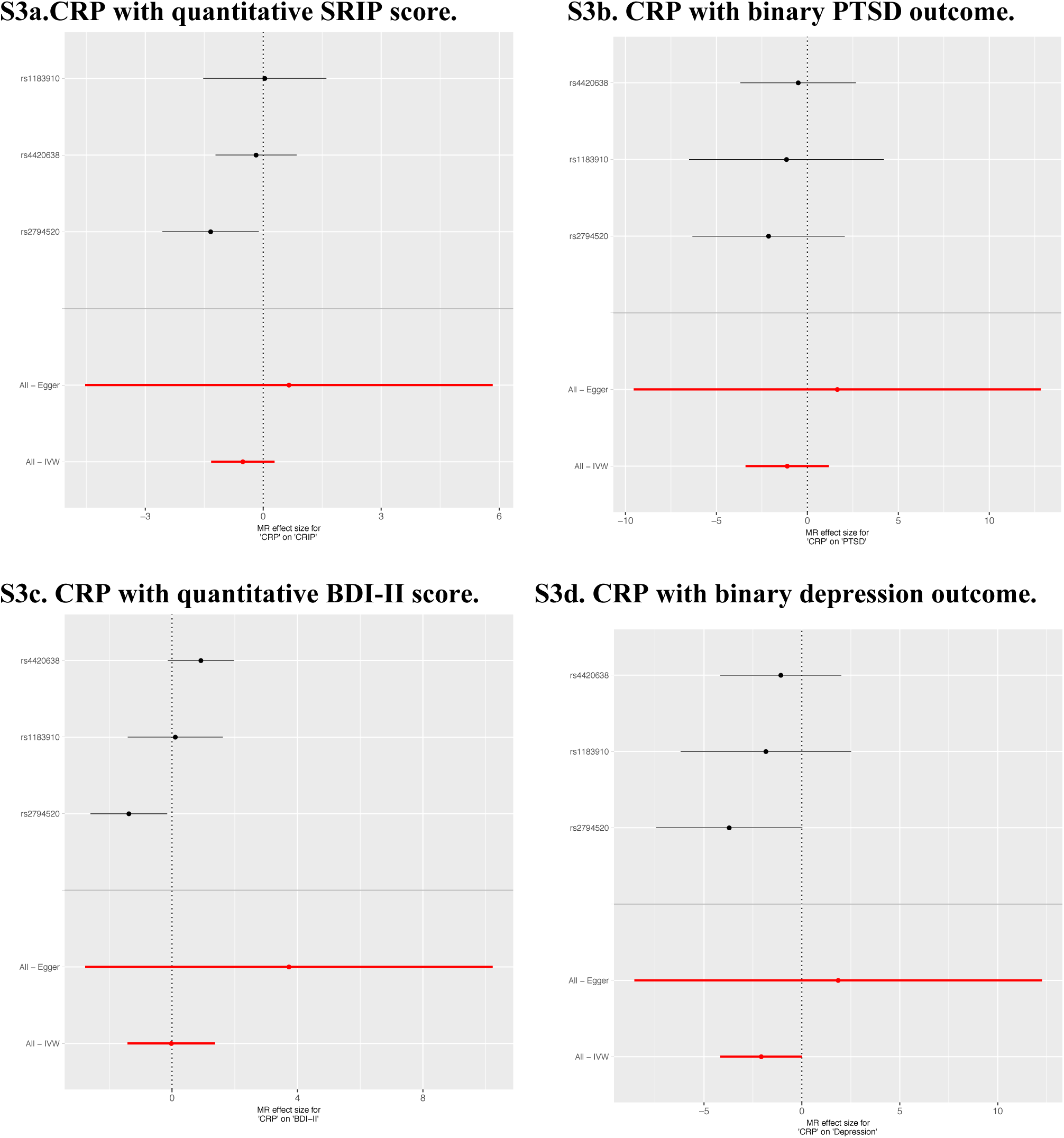
Meta-analysis forest plots (fixed-effects, inverse variance).

**Figure S4.**
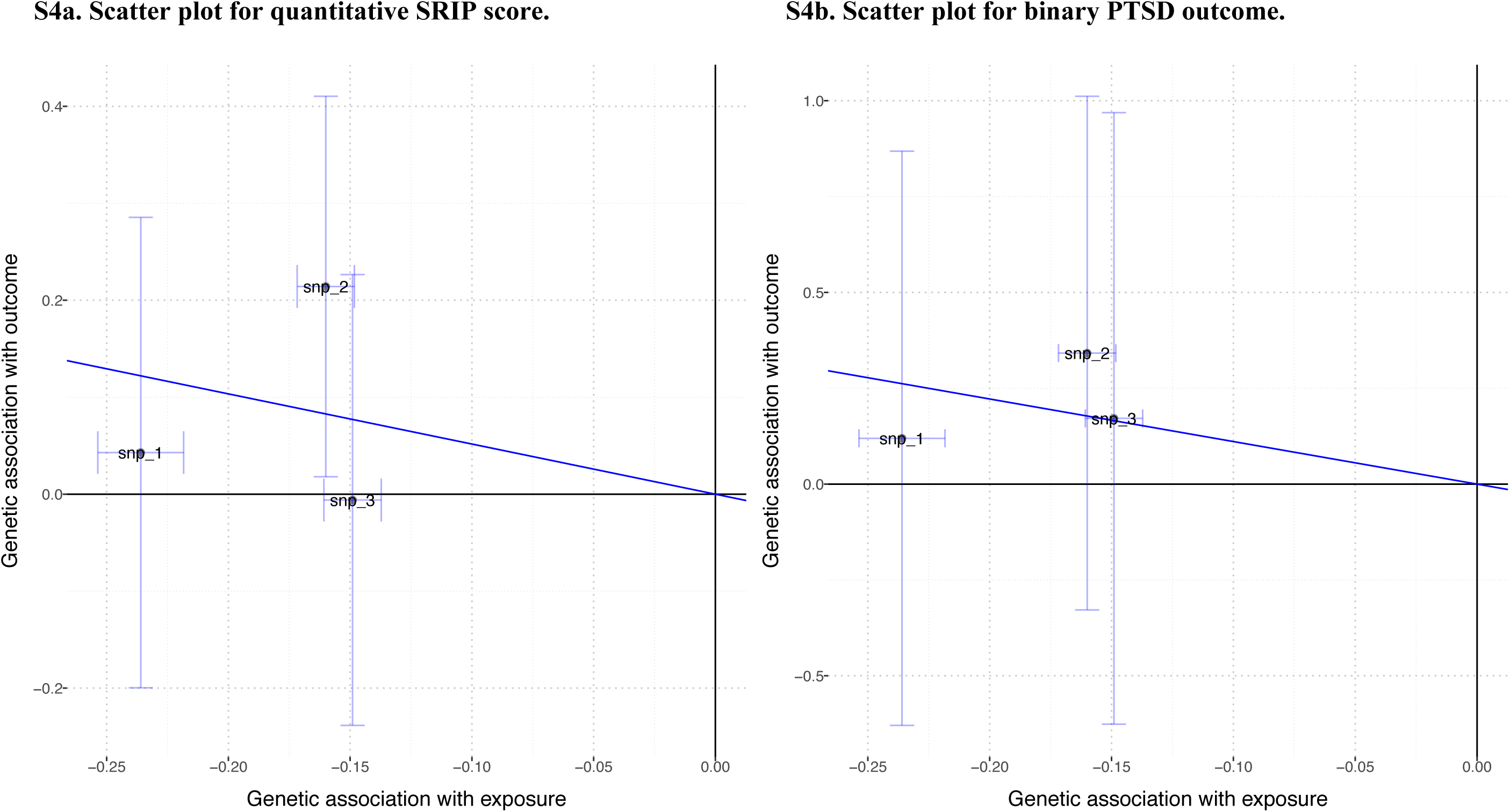
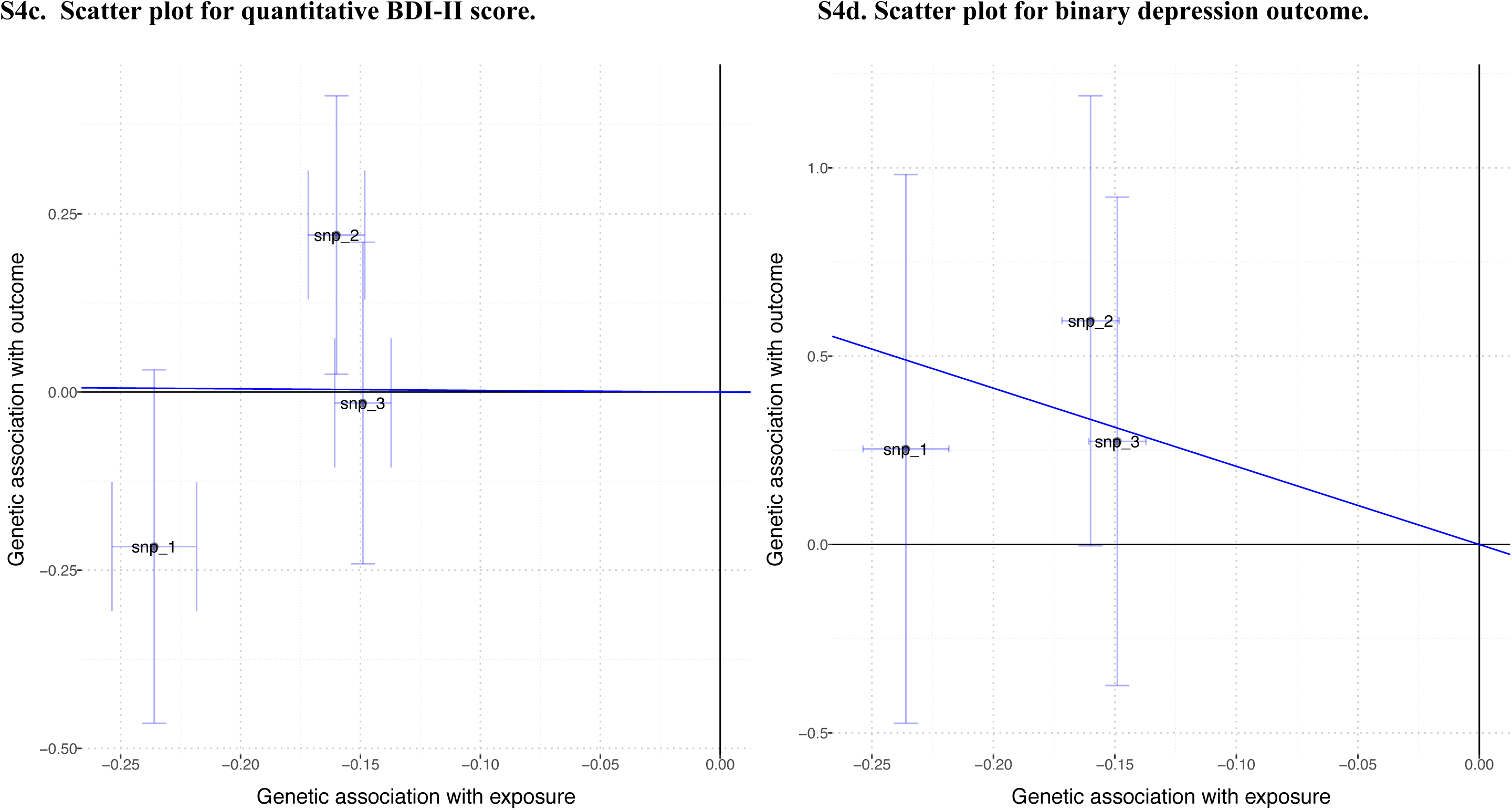
Scatter plots of associations between three SNPs, CRP, and psychopathology.

**Figure S5.**
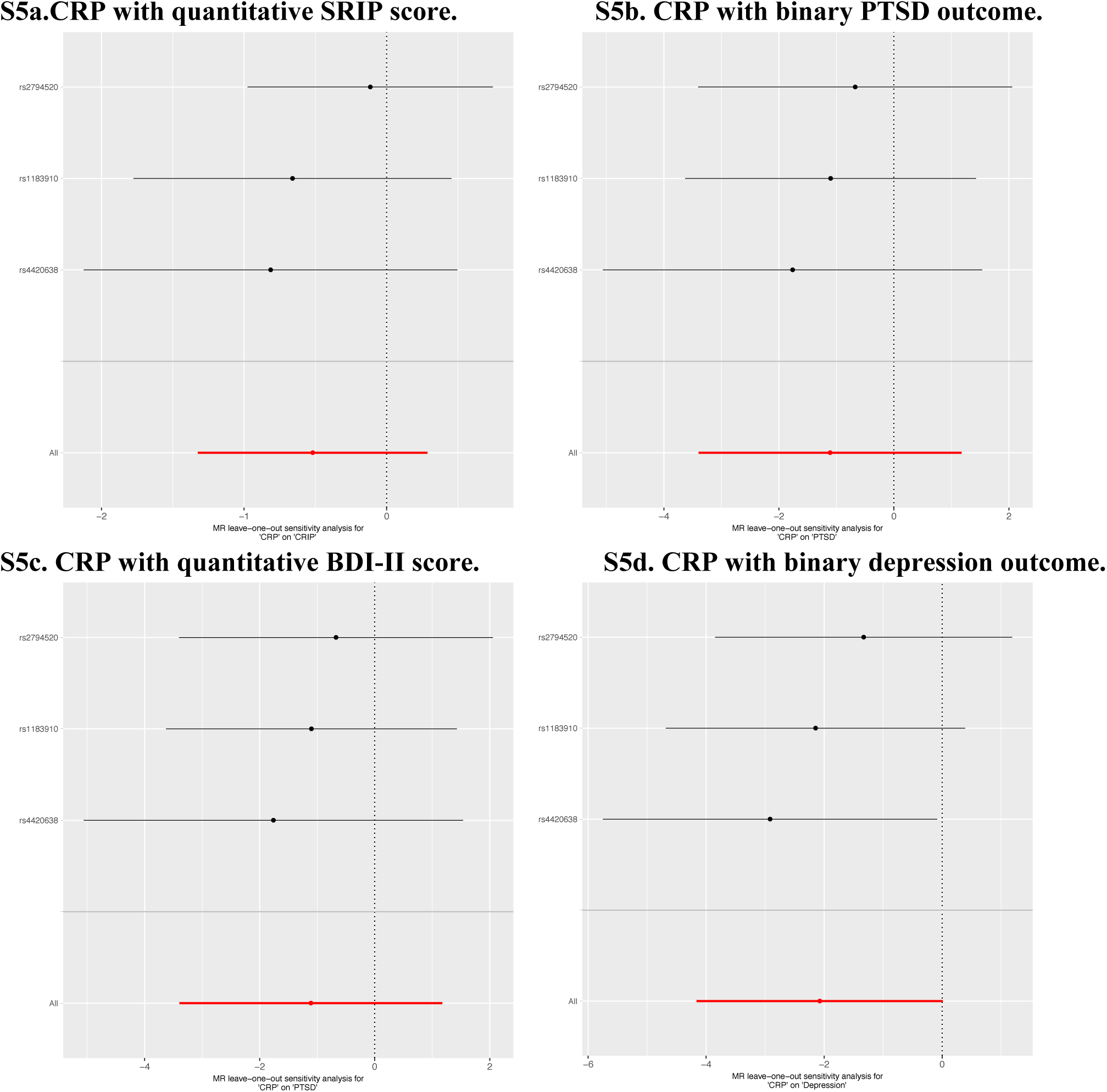
Forest plots of leave-one-out analysis.

**Table S1.** Genotyping information.

| SNP | Genotype | Distribution (N, frequency) | Minor | Observed | Expected |
| --- | --- | --- | --- | --- | --- |
|  |  |  | Allele | MAF | MAF |
| rs2794520 | TT/TC/CC | 129(0.12) / 491(0.45) / 464(0.43) | T | 0.3455 | 0.3421 |
| rs4420638 | GG/GA/AA | 48(0.05) / 313(0.29) / 710(0.66) | G | 0.1909 | 0.1510 |
| rs1183910 | AA/AG/GG | 94(0.09) / 451(0.41) / 537(0.50) | A | 0.2935 | 0.2955 |

**Table S2a.**
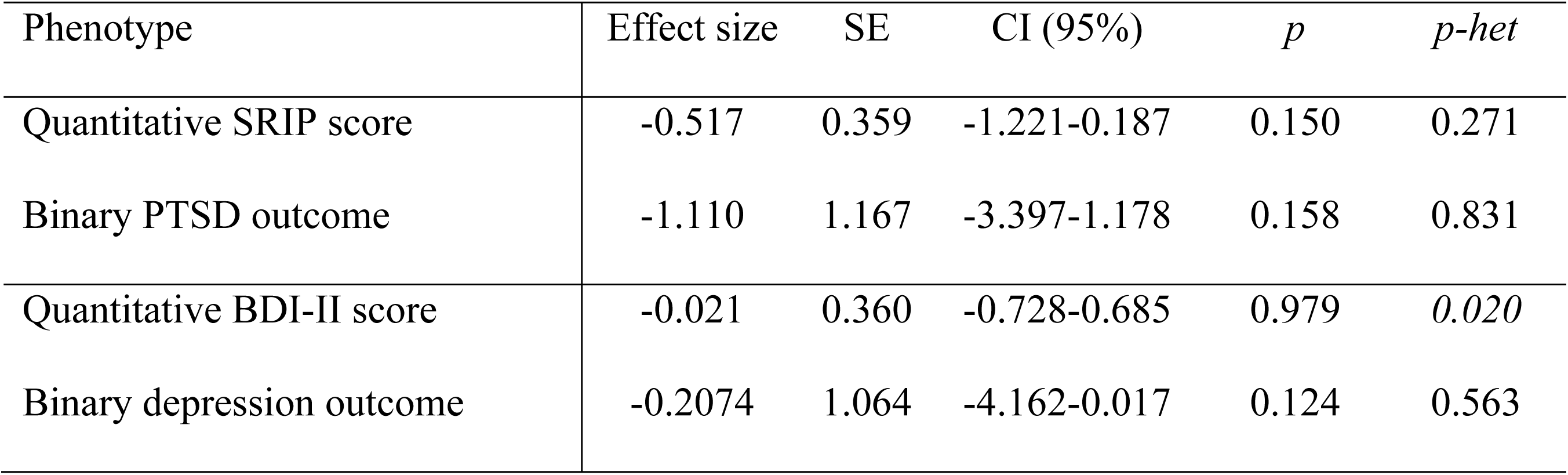
Meta-analysis summary statistics.

**Table S2b.**
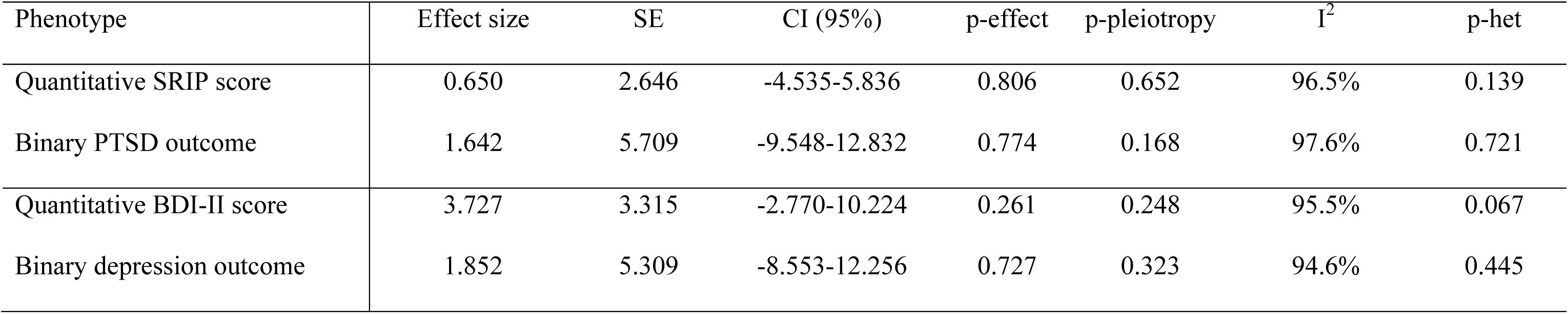
Meta-analysis summary statistics.

